# Detection of *Deformed wing virus* (DWV) in the Vietnamese Walking Stick *Medauroidea extradentata* (Phasmatodea)

**DOI:** 10.1101/2020.12.06.413948

**Authors:** Matan Shelomi, Wei Lin, Brian R Johnson, Michael J. Furlong, Kayvan Etebari

## Abstract

*Deformed wing virus* (DWV) is a single-stranded positive sense RNA virus which mainly infects honey bees (*Apis mellifera*) and can have devastating impacts on the colony. Recent studies have shown the presence of this virus in several species of *Apis* spp. and some other Hymenoptera, but our knowledge of their host range is very limited. We screened previously sequenced RNAseq libraries from different tissues of Vietnamese Walking Stick, *Medauroidea extradentata* (Phasmatodea) for DWV. We only found this virus in six libraries from anterior and posterior midgut tissue. From the midgut libraries we were able to construct the complete genome sequence of DWV, which consisted of 10,140 nucleotides and included one open reading frame. Pairwise genome comparison confirmed strong similarity (98.89%) of these assembled sequences with only 113 SNPs to the original DWV genome. Perhaps *M. extradentata* acquired this virus via a foodborne transmission by consuming DWV-infected material such as pollen or leaves contaminated with virus infected bee faeces.

---

*Deformed wing virus* (DWV; Picornavirales: *Iflaviridae*) is one of the most extensively studied viruses of honey bees, *Apis mellifera* (de Miranda and Genersch, 2010). DWV is a single-stranded, positive sense RNA virus with three master variants of varying pathogenicity (A, B, C) (Kevill et al., 2019). Its complete genome, with a single reading frame encoding a 328-kDa polyprotein, was published in 2006 (Lanzi et al., 2006). Its prevalence in *A. mellifera* colonies in some parts of the world can reach 100% (Williams et al., 2009), where it spreads horizontally through oral transmission (Mazzei et al., 2014) and vertically through infected semen and eggs (de Miranda and Fries, 2008). Such DWV infections are typically asymptomatic or “covert,” with the virus found in all bee tissues (de Miranda and Genersch, 2010). The parasitic Varroa mite (*Varroa destructor*, Varroidae, Parasitiformes) is a natural biological vector of five debilitating viruses including DWV and potentially 13 others. A recent study showed this parasite mite damage the host bees by consuming fat body (Ramsey et al., 2019). It has been shown that an “overt” DWV infection occurs when Varroa mite attacks the bees (Wilfert et al., 2016). These “overt” infections terminate in pupal or adult deformity and death, with devastating impacts on the colony (de Miranda and Genersch, 2010; Ramsey et al., 2019). Roberts et al., (2018) in a metagenomic study identified similar viruses to DWV in several honey bee populations across Australia in absence of its parasitic Varroa vector.

The high potential of RNA viruses to cross species barriers and become emerging infectious diseases (Schläppi et al., 2019) has led researchers to consider the extent of the host range of DWV. Using reverse transcription-polymerase chain reaction (RT-PCR) with specific primers, DWV has been found in at least four species of *Apis* (Zhang et al., 2012), eight other genera of Apidae, other Hymenoptera including ants (Formicidae) and wasps (Vespidae) (Martin and Brettell, 2019), and another bee mite, *Tropilaelaps mercedesae* (Mesostigmata, Laelapidae) (de Miranda and Genersch, 2010; Forsgren et al., 2009; Khongphinitbunjong et al., 2015). A study examining all arthropods inside established *A. mellifera* apiaries or within 800m of them found traces of DWV RNA in seven insect orders and two arachnid orders (Levitt et al., 2013). These included beekeeping pests like hive beetles and wax moths which are likely infected orally from bee food (Mazzei et al., 2014), and scavengers such as roaches and flies that were found under the hives and likely infected through infected material ejected from the hives. The study also found DWV in nearby plant-feeding beetles, bugs, and butterflies (Levitt et al., 2013); these were likely infected horizontally by feeding on pollen that had been contaminated by infected bees (Bailes et al., 2018; Mazzei et al., 2014). A recent study showed that DWV can replicate in a heterologous Lepidopteran cell line (P1) and harvested virus can also infect honey bees (Erez and Chejanovsky, 2020).

All present DWV studies are biased towards *A. mellifera* associated insects or other pollinators, and the RT-PCR primers involved were designed for DWV variants identified in *Apis*. This approach means that genetically diverse variants from non-bee hosts will be missed. However, next-generation sequencing (NGS) does not suffer from these structural constraints and it has the potential to reveal much of DWV’s true genetic and host diversity. As NGS data from insects that are not taxonomically, geographically, or behaviourally related to honey bees becomes increasingly available, the chances of finding DWV or related viruses in novel hosts grows. These findings “are of particular importance to determine the limits of host susceptibility, understand the frequency and mechanisms behind inter- and intraspecies transmission between non-*Apis* arthropods, and identify potential pathogenicity [of DWV in] non-managed insects” (Martin and Brettell, 2019).

Here we describe one such example, reporting on the host expansion of the virus into the Vietnamese stick insect *Medauroidea extradentata* (Phasmatodea: Phasmatidae). Native to the tropical forests of southern Vietnam, these easy to cultivate, wingless, parthenogenic, obligate leaf feeders are maintained internationally by researchers and Phasmatodea enthusiasts (Boucher and Varady-Szabo, 2005; Olive and Olive, 2019). Tolerant of subtropical and Mediterranean climates and a variety of host plants such as *Rubus* spp., *Rosa* sp., *Sorbus* spp. and *Fragaria ananassa*, the species has high invasion potential (Boucher and Varady-Szabo, 2005). The *M. extradentata* draft genome, Med v1.0 (GCA_003012365.1), was produced from such a wild-caught invader in California, USA (Brand et al., 2018). As the species is parthenogenic with few records of observed males, the genetics of the Californian and Vietnamese populations are expected to be highly similar.

In a previous study, the digestive tracts of three replicates of five laboratory-reared *M. extradentata* from the University of California-Davis, USA, were dissected and divided into anterior midgut, posterior midgut, ileum, rectum, head, and gut-free tissue of the rest of the body. To produce the host original transcriptome (Brand et al., 2018), RNA was extracted from each tissue type in TRIzol® (Invitrogen, USA) and quality was measured with an Agilent 2100 Bioanalyzer (Agilent, USA). Libraries were made using the Illumina TruSeq v2 kit (Illumina, Inc. San Diego, USA) and 100 bp paired-end sequencing performed on a HiSeq 4000 (University of California, Davis). The raw data was uploaded to NCBI, Accession Numbers SRR10437520-35.

In this study, the CLC Genomics Workbench version 12.0.1 (Qiagen, USA) was used for bioinformatics analyses. All libraries were trimmed from any remaining vector or adapter sequences. Low quality reads (quality score below 0.05) and reads with more than two ambiguous nucleotides were discarded. To identify novel viruses, all reads were mapped back to the *M. extradentata* draft genome (Brand et al., 2018) and unmapped reads were retained for *de novo* assembly. To check for possible cross contamination with honey bee RNA or presence of any potential varroa mite infestation, the reads were also mapped to the *Apis mellifera* Amel_HAv3.1 (GCA_003254395.2) and *Varroa destructor* (GCA_002443255.1) genome sequence. We couldn’t detect any sign of possible cross contamination or presence of *V. destructor* in the libraries.

The contigs were constructed with kmer size 45, bubble size 50, and minimum length of 500 bp, then corrected by mapping all reads against the assembled sequences (min. length fraction=0.9, maximum mismatches=2). The generated contigs were compared to the NCBI viral database using local BLAST and BLASTx algorithms. The e-value was set to 1×10-10 to maintain high sensitivity and a low false-positive rate. To detect highly divergent viruses, domain-based searches were performed by comparing the assembled contigs against the Conserved Domain Database (CDD) version 3.14 and Pfam v32 with an expected value threshold of 1×10−3. Sequences with positive hits to virus polymerase (RNA-dependent RNA polymerase (RdRp) domain: cd01699) were retained. The methods above were repeated for all other *M. extradentata* RNAseq data in GenBank (Table 1).

**Table 1:** *Medauroidea extradentata* transcriptomes in the NCBI database.

| Run Accession # | BioProject | Tissue | Location | DWV |
| --- | --- | --- | --- | --- |
| SRR2230573 | PRJNA286403 | head and thorax | Germany | No |
| SRR1172394 | PRJNA238833 | anterior midgut | USA: California | No |
| SRR1172402 | PRJNA238833 | posterior midgut | USA: California | No |
| SRR6383868 | PRJNA369247 | non-digestive tissue | USA: California | No |
| SRR6383869 | PRJNA369247 | non-digestive tissue | USA: California | No |
| SRR10437527 | PRJNA549703 | anterior midgut | USA: California | Yes |
| SRR10437534 | PRJNA549703 | anterior midgut | USA: California | Yes |
| SRR10437535 | PRJNA549703 | anterior midgut | USA: California | Yes |
| SRR10437531 | PRJNA549703 | ileum | USA: California | No |
| SRR10437532 | PRJNA549703 | ileum | USA: California | No |
| SRR10437533 | PRJNA549703 | ileum | USA: California | No |
| SRR10437520 | PRJNA549703 | non-digestive tissue | USA: California | No |
| SRR10437521 | PRJNA549703 | non-digestive tissue | USA: California | No |
| SRR10437522 | PRJNA549703 | non-digestive tissue | USA: California | No |
| SRR10437523 | PRJNA549703 | non-digestive tissue | USA: California | No |
| SRR10437524 | PRJNA549703 | posterior midgut | USA: California | Yes |
| SRR10437525 | PRJNA549703 | posterior midgut | USA: California | Yes |
| SRR10437526 | PRJNA549703 | posterior midgut | USA: California | Yes |
| SRR10437528 | PRJNA549703 | rectum | USA: California | No |
| SRR10437529 | PRJNA549703 | rectum | USA: California | No |
| SRR10437530 | PRJNA549703 | rectum | USA: California | No |

The identified DWV genome in *M. extradentata* was built out of 71,113 assembled reads from six RNAseq libraries from the anterior and posterior midgut tissue. No reads associated with DWV were found in other tissues from this transcriptome set, or from any other *M. extradentata* SRA data in GenBank (Table 1). Over 98.9% of reads from the *M. extradentata* transcriptomes were mapped to the *M. extradentata* genome, and no evidence of RNA contamination with another insect or organism was detected. We conclude that the transcript represents a true DWV, and the annotated genomic sequence of this virus has been deposited in GenBank under the accession number MW222481.

Only around 0.02% of the total reads in these RNAseq libraries mapped to DWV genome, which is similar to what is found in mosquito midguts during an active Dengue infection (Bonizzoni et al., 2012). The pattern of mapping showed higher coverage towards the RdRp region and 3’ ends, which is probably due to bias in reverse transcription of the oligo dT primer (Figure 1).

**Figure 1.**
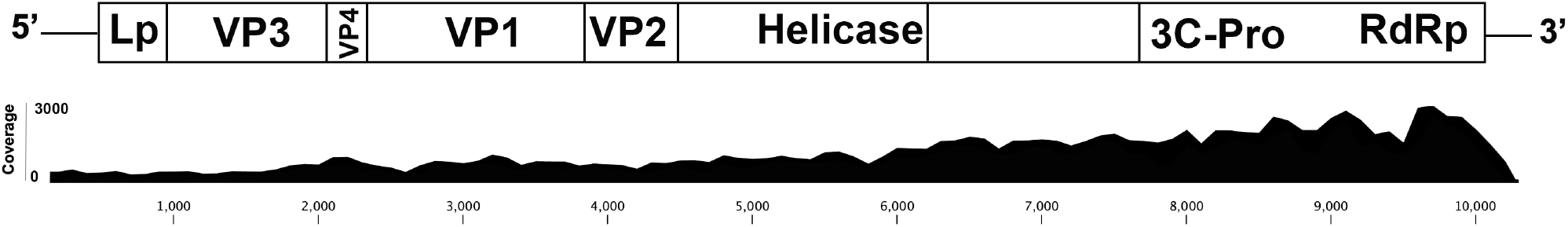
Schematic presentation of the genome of DWV identified in *Medauroidea extradentata*. The location of predicted conserved domains is in the upper panel. The reads were mapped back to the sequenced genome to display the coverage and sequence depth (lower panel).

A single polyprotein structure with several conserved domains such as Picornavirus capsid protein (Rhv-like), RNA helicase, Peptidase and RNA-dependent RNA polymerase (RdRp) was predicted, as this is consistent with the genome arrangement of other DWVs. Once the viral sequences were confirmed as DWV, the deduced amino acid sequence of the complete polypeptide for several DWVs obtained from GenBank were used with the consensus DWV sequence obtained from the *M. extradentata* to create a maximum likelihood phylogeny tree (Figure 2). Protein distance was measured by the Jukes-Cantor algorithm to construct the tree topologies and its statistical significance was calculated based on 1000 bootstraps.

**Figure 2.**
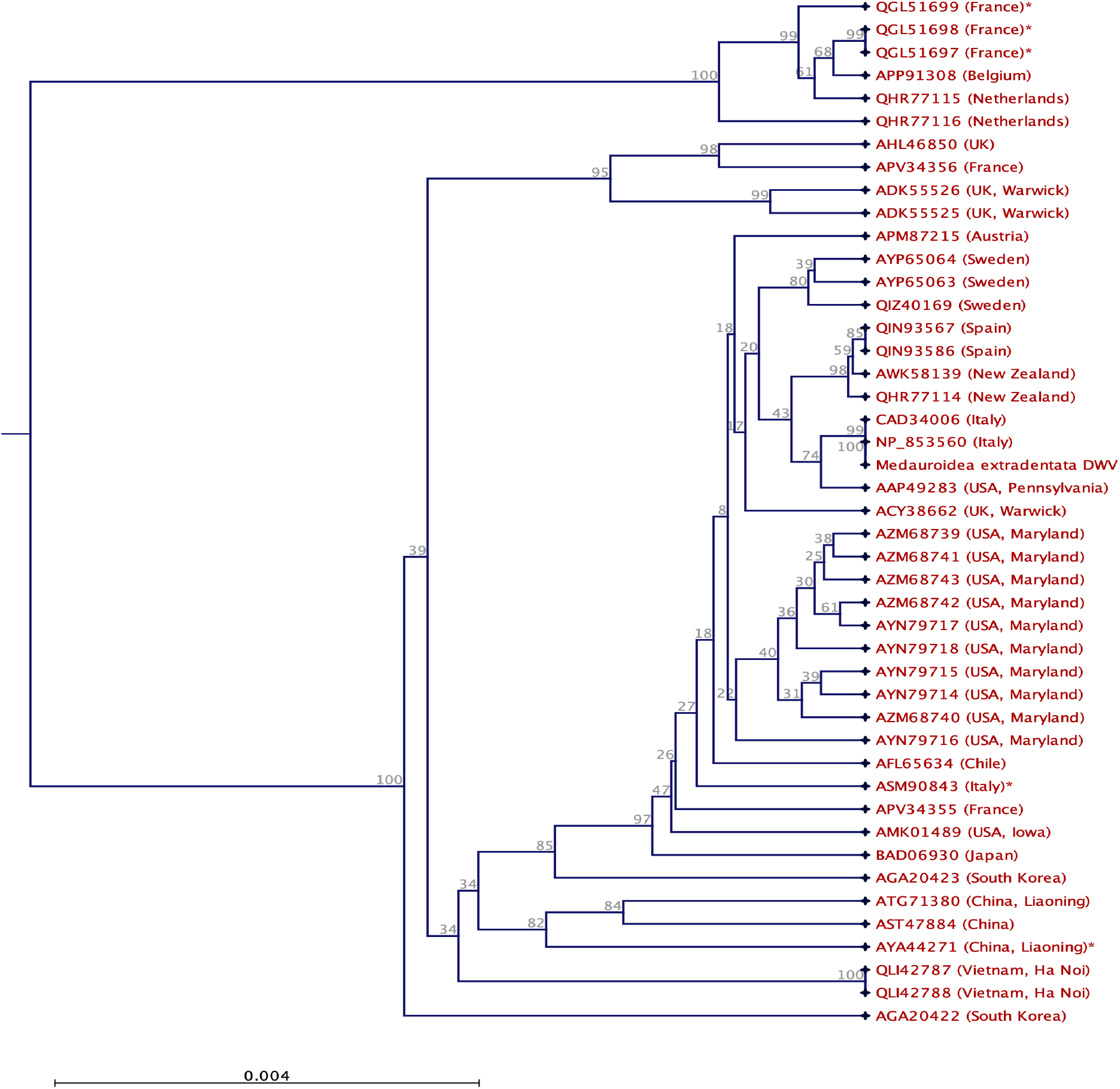
Maximum likelihood phylogeny based on the deduced amino acid sequence of DWV Polyprotein. Based on sequence similarity, the virus identified from *Medauroidea extradentata* anterior and posterior midgut tissue is unmistakably a strain of DWV, and likely from the DWV-A variant group. Protein distance was measured by the Jukes-Cantor algorithm to construct the tree topologies and its statistical significance was calculated based on 1000 bootstraps.

Pairwise genome comparison confirmed strong similarity (98.89%) of these assembled sequences with only 113 SNPs to the original DWV genome (Accession Numbers NP_853560, CAD34006.2) (Lanzi et al., 2006). The maximum likelihood phylogeny (Figure 2) also placed the newly identified *M. extradentata* virus firmly in with European-American strains of DWV. The phylogenic analysis of the deduced amino acid sequence of the polyprotein in different geographical isolates of DWV separated some European DWV isolates from others. The Asian isolates of DWV are also separated from other groups. However, some of those DWV were isolated from different species of *Apis* and this might have an impact on their polyprotein sequence structure. Based on amino acid sequence analysis, this newly identified DWV likely belongs to the variant group DWV-A.

This is the first study to report presence of DWV in the *M. extradentata* transcriptome which is an evidence of DWV spill over in a Phasmatodea. No DWV were found in any other published *M. extradentata* transcriptomes, including those made from RNA extracted from other individuals from the same institution (Shelomi et al., 2014), and an RNASeq study of a whole juvenile from the same study (Brand et al., 2018). DWV is thus not found in 100% of *M. extradentata*. The phasmids in this study (Brand et al., 2018) were fed rose (*Rosa* sp.) leaves, and so could have consumed DWV-infected material such as pollen that had fallen onto the leaves. Whether there are other ways for insects to become infected with DWV independent of bee-related plant parts is unknown. Recent studies have experimentally identified the role of shared flowers and other plant materials in the transmission of some of bee pathogens (Alger et al., 2019; Figueroa et al., 2019). Yañez et al., (2020) reviewed the possible routes of transmission of bee viruses into other arthropods and concluded that the oral-fecal transmission is most likely the main route of virus transmission into non-bee spices (Yañez et al., 2020).

Positive-sense ssRNA viruses like DWV do not require transcription, but function as mRNA themselves; so the presence of DWV RNA in a transcriptome does not necessarily indicate active replication, but merely the presence of the virus. Here, we recovered more than 70,000 DWV associated reads from six RNAseq libraries, which rules out accidental contamination during sample processing and sequencing. To rule out passive acquisition of DWV during feeding, we checked for the presence of other bee viruses such as Sacbrood virus (AF092924), Black queen cell virus (BQCV: AF183905), Acute bee paralysis virus (AF150629), Lake Sinai virus (KX883223), Kashmir bee virus (AY275710), Israeli acute paralysis virus (EF219380) and Chronic bee paralysis virus in these libraries. It has been demonstrated that some other honey bee viruses, particularly BQCV, maintain high prevalence alongside DWV in infected bees (Chen et al., 2004; Roberts et al., 2018). We could not detect any significant number of reads associated to any other bee viruses in DWV infected *M. extradentata* midgut tissues (Table 2). We only found 196 and 32 reads associated with Sacbrood virus and BQCV respectively, compared to over 70,000 for DWV.

**Table 2.**
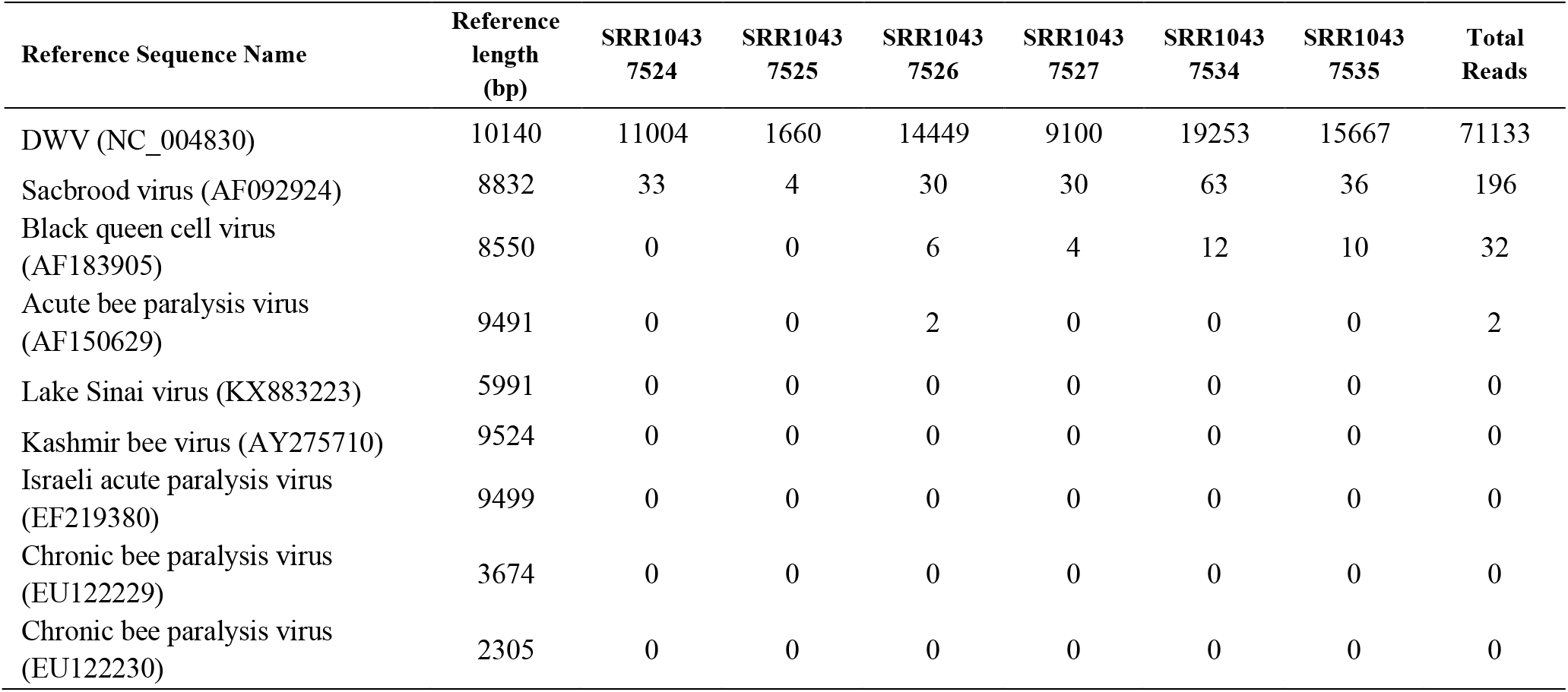
number of reads mapped to common honey bee’s viruses

| Reference Sequence Name | Reference length (bp) | SRR1043 7524 | SRR1043 7525 | SRR1043 7526 | SRR1043 7527 | SRR1043 7534 | SRR1043 7535 | Total Reads |
| --- | --- | --- | --- | --- | --- | --- | --- | --- |
| DWV (NC_004830) | 10140 | 11004 | 1660 | 14449 | 9100 | 19253 | 15667 | 71133 |
| Sacbrood virus (AF092924) | 8832 | 33 | 4 | 30 | 30 | 63 | 36 | 196 |
| Black queen cell virus (AF183905) | 8550 | 0 | 0 | 6 | 4 | 12 | 10 | 32 |
| Acute bee paralysis virus (AF150629) | 9491 | 0 | 0 | 2 | 0 | 0 | 0 | 2 |
| Lake Sinai virus (KX883223) | 5991 | 0 | 0 | 0 | 0 | 0 | 0 | 0 |
| Kashmir bee virus (AY275710) | 9524 | 0 | 0 | 0 | 0 | 0 | 0 | 0 |
| Israeli acute paralysis virus (EF219380) | 9499 | 0 | 0 | 0 | 0 | 0 | 0 | 0 |
| Chronic bee paralysis virus (EU122229) | 3674 | 0 | 0 | 0 | 0 | 0 | 0 | 0 |
| Chronic bee paralysis virus (EU122230) | 2305 | 0 | 0 | 0 | 0 | 0 | 0 | 0 |

This finding suggests that *M. extradentata* is a host for DWV and that the sequencing wasn’t just picking up DWV particles in the gut contents. Although the source of virus could have been food material infected with bee faeces, probably only DWV could replicate inside the *M. extradentata* midgut, and other bee viruses could not replicate and produce progeny virions. The *M. extradentata* had no symptoms of acute, overt DWV infection as found in *Apis mellifera* infected through varroosis. DWV infection in *M. extradentata* appears to be limited to the midgut and would thus be a persistent covert infection, while in *Apis mellifera* DWV replicates in all tissues as a covert or overt infection (de Miranda and Genersch, 2010).

The geographic distribution of DWV in *M. extradentata* and whether it exists in individuals in the species’ native habitat of Southeast Asia, where DWV also originated, is also unknown. Comprehension of DWV transmission strategies among different developmental stages and tissues of *M. extradentata* is necessary to explain the biological and ecological importance of this virus, which appears to be non-pathogenic. Further research is still needed to understand the role of different host species for this virus.

## Statement of Author Contributions

**Matan Shelomi:** Conceptualization, Methodology, Investigation, Writing-Original draft preparation.

**Wei Lin:** Investigation, Resources

**Brian R Johnson:** Investigation, Project administration, Funding acquisition, Writing-Reviewing and Editing

**Michael J. Furlong:** Investigation, Project administration, Funding acquisition, Writing-Reviewing and Editing

**Kayvan Etebari:** Conceptualization, Methodology, Investigation, Data curation, Visualization, Project administration, Writing-Reviewing and Editing

